# Rhythm generating mechanisms in rat sino-atrial node and ventricle

**DOI:** 10.1101/2023.02.28.529494

**Authors:** Jesi Charles, Latha Nedumaran, Swetha Raman, Elizabeth Vinod, Rajalakshmi Rajasegaran, Kamalakannan Vadivel, Anand Bhaskar, Sathya Subramani

**Affiliations:** Department of Physiology, Christian Medical College, Vellore

## Abstract

The major membrane currents responsible for sinoatrial and idioventricular rhythm-generation were studied in isolated rat heart preparations, perfused in Langendorff mode. The rates of whole isolated hearts beating with sinoatrial rhythm decreased with cesium and ivabradine, both blockers of the funny current, and were not affected by nickel, at a dose which blocks T-type calcium current. The sinoatrial rhythm was completely abolished by reduction or removal of sodium from the perfusate (interventions that inhibit calcium-extrusive mode of the sodium/calcium exchanger), or by nifedipine, an L-type calcium channel blocker. Idioventricular rhythm, however, was arrested only by reduction of sodium in the perfusate. Ivabradine reduced the idioventricular rate, nickel did not cause any change, while nifedipine in some cases increased it. The inferences made based on these observations are that INCX and ICaL are obligatory rhythm-generating currents in the sinoatrial node, while INCX is the only obligatory mechanism for an idioventricular rhythm. The funny current is not an obligatory requirement for sinoatrial as well as idioventricular rhythm-generation. However, it enhances the frequency of LCRs. Our results in the isolated whole heart are in corroboration with results from isolated cells.

## Introduction

In mammalian heart, automaticity is initiated and maintained by the sino-atrial node (SA node), which is a group of specialized cells generating spontaneous action potentials rhythmically (Verheijck et al., 1998; Bouman and Jongsma, 1986). The ability to generate spontaneous action potentials is not restricted to the SA node (Meijler and Janse, 1988; Callewaert et al., 1μ984). However, it behaves as the primary pacemaker because the rate of generation of action potentials is highest in this region. The SA node action potential consists of a phase of diastolic depolarization (phase 4), followed by the depolarization phase (phase 0) which is then followed by repolarization (phase 3). Phase 4, which is also called as ‘pacemaker potential’ or ‘spontaneous diastolic depolarization’ is responsible for automaticity. Presently, it is thought that the spontaneous diastolic depolarization is primarily due to activation of the following inward currents: *Hyperpolarization activated funny current* (I_f_), *Voltage gated T-type* (I_CaT_) and *L-type calcium currents* (I_CaL_) *and current through Sodium /Calcium Exchanger* (I_NCX_) (Difrancesco and Zaza, 1992; Maltsev et al., 2006; Mangoni et al., 2003; Huser et al., 2000). The probable sequence of these currents, as we understand is what underlies the perceived dichotomy between the “membrane clock” and the “calcium clock” theories of rhythm-generation. These theories are discussed below.

Subsequent to an SA node action potential, membrane repolarization by outward K^+^ currents, takes the membrane to its ‘Maximum Diastolic Potential’ (MDP). At such relatively hyperpolarized potentials negative to −45 mV, the funny current If, is activated. If is proposed to initiate diastolic depolarization (Difrancesco 2010, Mangoni et al, 2008). T-type calcium channels are low-voltage activated and are also open at voltages negative to −40 mV (Huser et al 2000). They are suggested to be responsible for late phase diastolic depolarization. Sarcoplasmic reticulum releases calcium (Diastolic calcium release or local Ca^2+^ release (LCR) in current terminology) during diastole ((Huser et al., 2000). Efflux of this calcium via NCX (forward mode) generates an inward current, given that the stoichiometry of NCX is 3 Na^+^: 1 Ca^++^ (Kimura et al., 1987). Inward current through NCX (I_NCX_) is now recognized as a predominant pace-maker mechanism (Maltsev & Lakatta, 2007) responsible for diastolic depolarization. It is suggested that Ca^2+^ influx through T type Ca^2+^ channels is what triggers the LCR through ryanodine receptors (RYR) (Huser et al., 2000), which then drives forward mode NCX, but this view is debatable (Li et al, 2011, Hagiwara et al 1988). The LCR-dependent inward current by NCX will take the membrane to a more positive potential at which the L-type calcium channels will be activated, and are responsible for the last phase of diastolic depolarization (Mangoni 2003) and the upstroke of the action potential of the sinus node (Hagiwara et al, 1988). The above mechanisms can be considered as the “Membrane clock” theory where depolarizing events are initiated primarily at the cell membrane, starting with If. The alternative theory is that of a “Calcium clock”, which hypothesizes that diastolic calcium release from SR (LCR) (extrusion of this calcium through NCX being the primary depolarizing mechanism) is spontaneous and rhythmic and is not dependent on membrane depolarization (Maltsev and Lakatta, 2007). What we understand as the basic difference between the roles of NCX in “membrane clock” and the “calcium clock” theories is as follows: The membrane clock theory places emphasis on the role of If and also suggests that calcium release from SR to drive NCX in forward mode is triggered by membrane depolarization and/or calcium entry through T-type calcium channels. On the other hand, the calcium clock theory suggests that the release of calcium from SR is spontaneous and is not dependent on events at the cell membrane. That diastolic SR calcium release is spontaneous and can be independent of cyclical voltage changes in the membrane has been demonstrated (Maltsev and Lakatta, 2007). However, a dynamic interaction between the membrane and calcium clocks has been repeatedly emphasized, the two clocks being entrained during rhythm-generation (Maltsev and Lakatta, 2008). The details of such entrainment are however unclear. Possibly, hyperpolarizing potentials enhance the probability of LCR, in addition to activating I_f_. A good description of the two theories, and the more recent coupled-clock theory are found in Mesirca *et al* (2015).

The contributions of the currents mentioned above to cardiac rhythm generation have been studied mostly at the cellular level. There is, however, a paucity of studies on the whole isolated heart. Here we report results of experiments conducted to assess the contribution of the four stated inward membrane currents towards rhythm generation in two different preparations namely, the isolated rat heart with SA nodal rhythm and isolated rat heart with idio-ventricular rhythm. The results may provide some answers to the debate (Difrancesco 2010; Maltsev et al., 2006) on the extent of contribution to pace-making of the funny current (the membrane clock) versus the inward current through NCX triggered by spontaneous LCR (calcium clock). The isolated heart preparation is also a useful preparation for screening of toxins and drugs for effects on cardiac rhythm, especially, putative anti-arrhythmic agents.

## Materials and methods

### Ethical approval

The experiments were approved by the Institutional Animal Ethics Committee of Christian Medical College, Vellore, India. (IAEC Nos: 3/2012, 5/2012, 9/2012 and 14/2012). Wistar rats (*Rattus norvegicus*) of both sexes, weighing between 200 to 300 g were used.

### Anaesthesia

The rats were anesthetized with intraperitoneal Ketamine at a dose of 100mg/Kg body weight.

### Experimental Protocol

After deep anaesthesia, a sub-costal incision was made. The diaphragm was cut, and chest-cavity was opened. Hearts were isolated and perfused with Physiological salt solution (PSS) with a cannula placed in the aorta above the aortic valve, to perfuse the coronary arteries (Langendorff preparation). For experiments on idio-ventricular rhythm the right atrium was carefully excised after cannulation and the atrio-ventricular bundle was severed by a cut in the inter-ventricular septum (Benforado, 1958). ECG electrodes in contact with the surface of heart, were used to record the surface electrogram. Signals were acquired and digitized by a computerized data acquisition system (CMCdaq). Heart rate was calculated at different time points from the recorded electrogram, by counting the R waves.

The isolated hearts were perfused with the Physiological Salt Solution (PSS) for at least 15 minutes or till the heart rate stabilized. Then the perfusate was changed to the test solution which was perfused for 15 minutes or till the heart rate stabilized at a new level after the intervention. This was followed by a wash step – perfusing with PSS for another 15 minutes.

## Solutions used

The ‘*Physiological Salt Solution’* (PSS) contained (in mmol/L): 135 NaCl, 5.4 KCl, 0.4 NaH_2_PO_4_, 3 MgCl_2_, 1 CaCl_2_, 10 HEPES, 10 Glucose. pH was 7.4 titrated with 1 mol/L NaOH.

The *‘low sodium solution’* contained (in mmol/L): 35 NaCl, 100 choline chloride, with other compositions like PSS and the pH was titrated to 7.4 with 1 mol/L NaOH.

The *‘sodium-free solution’* contained (in mmol/L): 135 LiCl instead of NaCl. Other constituents were the same as PSS. The pH was titrated to 7.4 with 1 mol/L LiOH.

The *‘high calcium solution’* contained (in mmol/L) 5 CaCl_2_ and other constituents were the same as PSS. The pH was 7.4 and titrated with 1 mol/L NaOH.

### Drugs and chemicals

Ivabradine was used at a concentration of 10 μmol/L to block the funny current. Cesium (5 mmol/L) was also used as a blocker of the funny current. Two different interventions were performed to inhibit the inward current due to the sodium-calcium exchanger - these were (a) low sodium solution (b) sodium-free solution containing Lithium. L-type calcium current was inhibited with Nifedipine (10 μmol/L). Nickel (40 μmol/L) was used to block T-type calcium channels. Cesium chloride, Lithium hydroxide and Lithium chloride were purchased from Lobachemie. Ivabradine, Nifedipine, CaCl2, MgCl2 and glucose were purchased from Sigma-Aldrich. Disodium hydrogen phosphate was purchased from Qualigens. NaOH and Nickel chloride were purchased from Merck.

## Analysis

Wilcoxon’s signed rank test was used to compare pre intervention heart rate with heart rate after intervention when it had stabilized. P values <0.05 were considered significant. For graphic depiction, post-intervention heart rates were normalized to the pre-intervention level. Mean and SD were calculated for normalized post-intervention heart rates at different time points from 6 experiments and graphs were generated using IGOR Pro 5.0 software. Mean and SD values for heart rates were corrected to the nearest whole number.

## Results

### Whole heart experiments to assess SA nodal rhythm generation

The basal heart rate with PSS, averaged from pre-intervention values of all experiments was 117 ± 29 beats per minute (bpm), n= 40.

### Effects of Cesium

The basal heart rate with PSS in this set of experiments was 120 ± 25 bpm. When 5 mmol/L cesium chloride was added to the perfusate, heart rate reduced to 87 ± 16 bpm after 5 minutes and remained at around that value for the rest of the period when cesium chloride was present in the perfusate. The reduction was statistically significant. (p=0.027, n=6). Following wash with PSS, rate revived to near normal levels (107 ± 25 bpm). When post-intervention values were normalized to pre-intervention heart rates, there was a reduction of 27 ± 5% after 5 minutes of cesium perfusion (Figure 1)

**Figure1:**
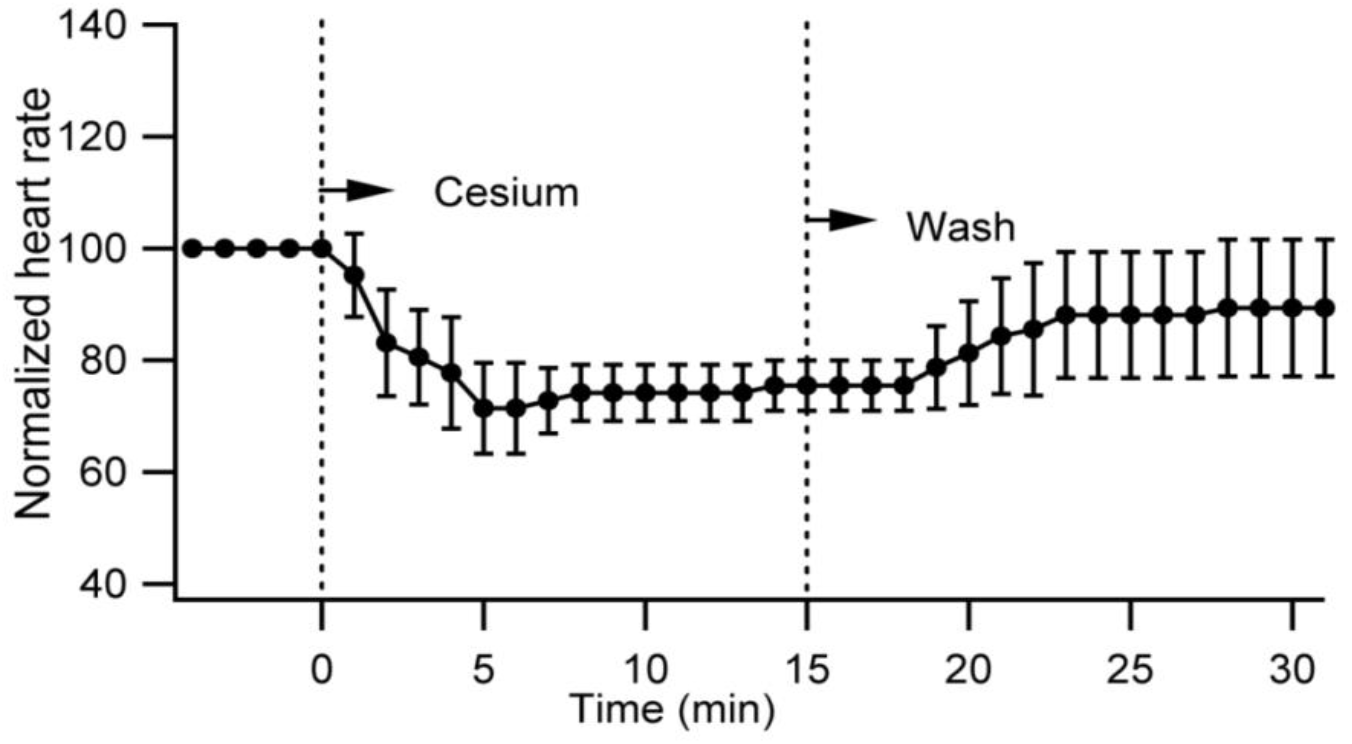
Cesium (5 mmol/L) which blocks If, reduced heart rate, but did not stop the heart in isolated perfused rat hearts (n = 6)

### Effects of Ivabradine

With Ivabradine (10 μmol/L), the heart rate reduced from 136 ± 36 to 74 ± 29 bpm. (p = 0.043, n = 5) With normalization of post-intervention heart rates as percentages of the pre-intervention value, there was a 47 ± 13% reduction in heart rate during intervention. There was no recovery when Ivabradine was stopped, and the heart was perfused with PSS (the ‘wash’ step). (Figure 2)

**Figure 2:**
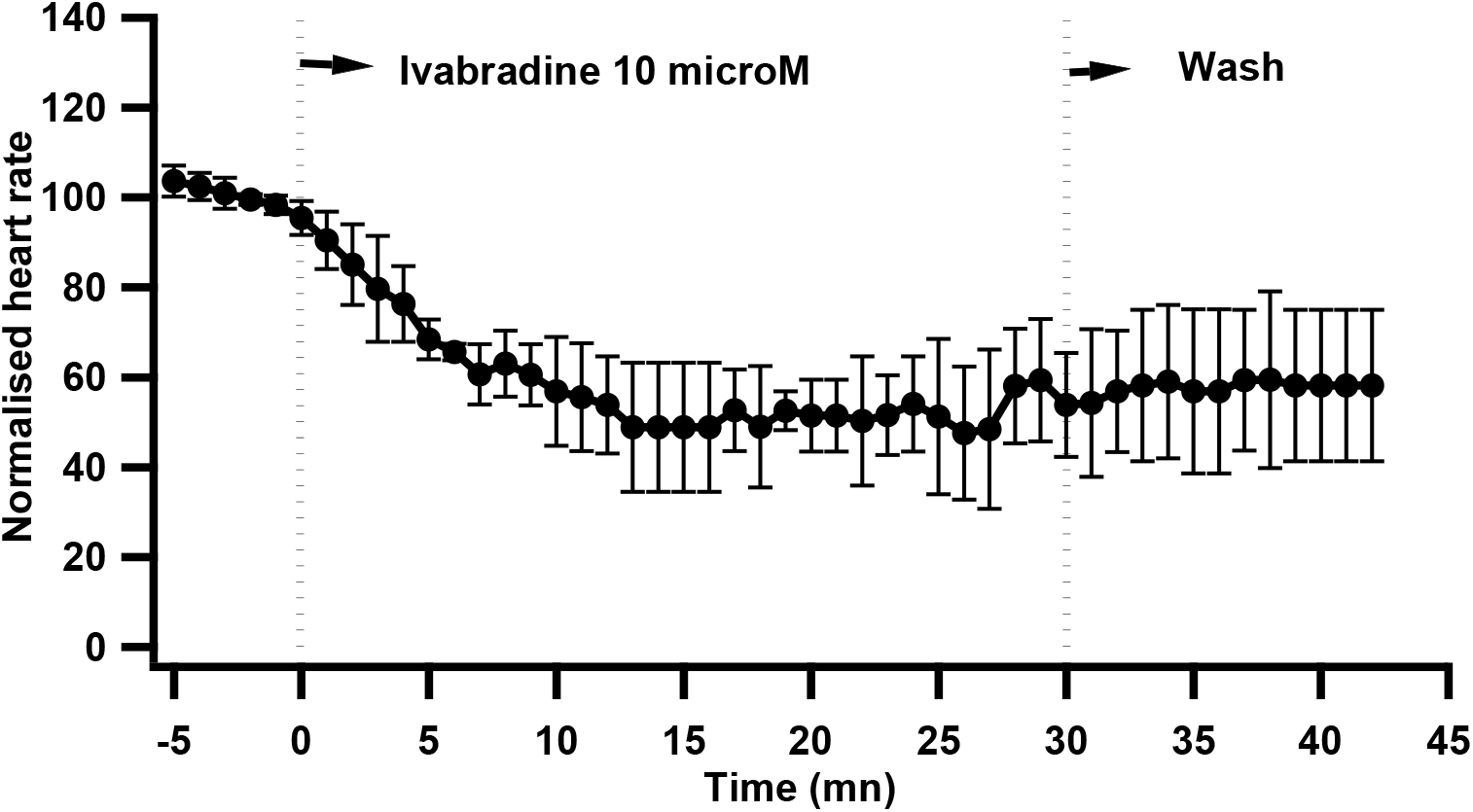
Ivabradine (10 μmol/L) which blocks If, reduced the heart rate in isolated perfused rat heart. It however did not stop the heart (n =5)

### Effects of Low sodium

Isolated hearts with a basal heart rate of 122 ± 28 bpm in PSS, progressively slowed down and stopped beating by 10 minutes (p value = 0.027, n=6), upon switching to a low sodium solution. After sodium replacement by perfusing with PSS, heart beat slowly recovered (79 ± 33 bpm) but did not reach pre-intervention values even after 20 minutes of perfusion with PSS (Figure 3).

**Figure 3:**
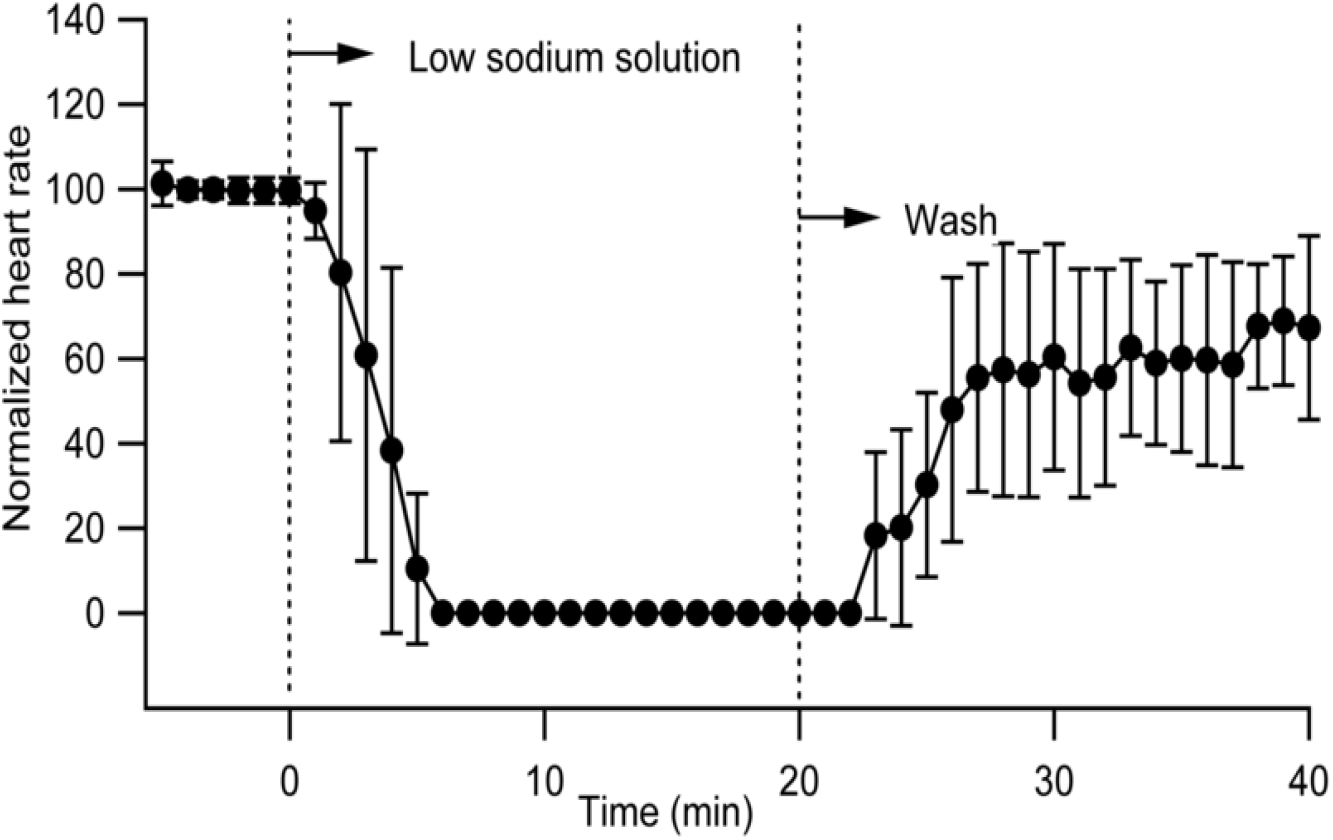
Low sodium solution (35 mmol/L sodium) stopped the isolated rat heart within 10 minutes of perfusion.

### Effects of Sodium-free Lithium extracellular solution

With sodium free solution (sodium chloride replaced with lithium chloride as lithium can carry current through voltage gated Na^+^ channel while it cannot substitute for Na^+^ on the NCX), heart stopped completely by the 12^th^ minute of change to lithium ECF (p value=0.028) The heart rate recovered to 67 ± 36 % of the pre-intervention value by 30 minutes after wash with PSS (Figure 4).

**Figure 4:**
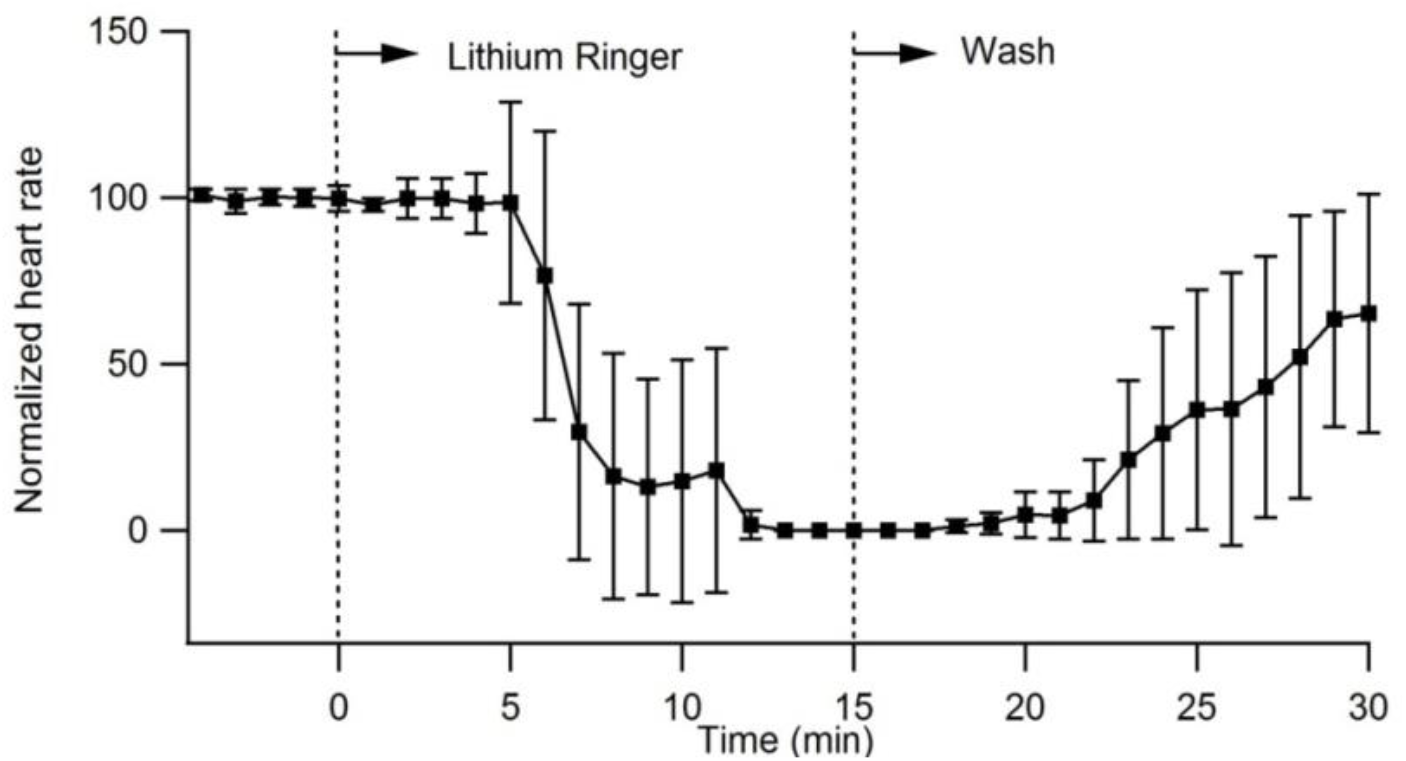
Sodium-free Lithium solution stopped the isolated rat heart within 12 minutes of perfusion. (n = 6)

### Effects of high calcium solution (5 mmol/L Calcium)

High calcium solution increased the heart rate in 3 experiments while there was a slight decrease in 3 other experiments. Representative raw tracings are shown in Figure 5. The change in heart rate with high calcium is therefore concluded as equivocal.

**Figure 5:**
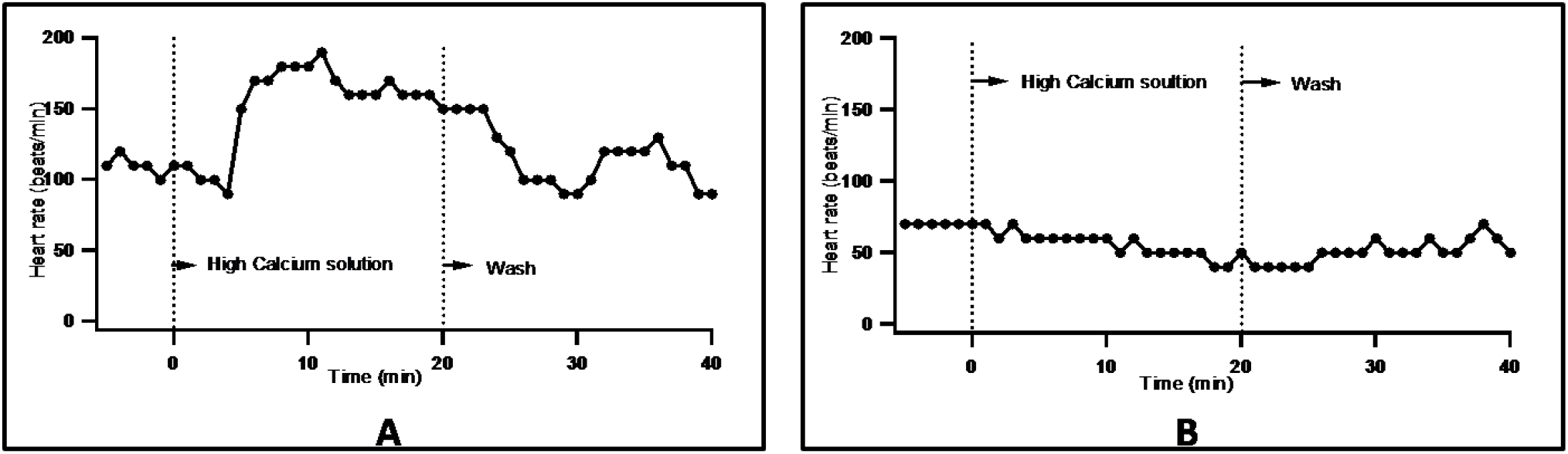
A, B: High calcium in the perfusate produced equivocal responses, increasing the rate in 3 experiments and decreasing the rate in 3 other experiments.

### Effects of Nifedipine

With Nifedipine (10 μmol/L), the heart stopped completely between 10 to 15 minutes of intervention (Figure 6). The basal heart rate before intervention was 109 ± 9 bpm. The 100% blockade of heart rate with Nifedipine was statistically significant with a p value of 0.042 (n = 5).

**Figure 6:**
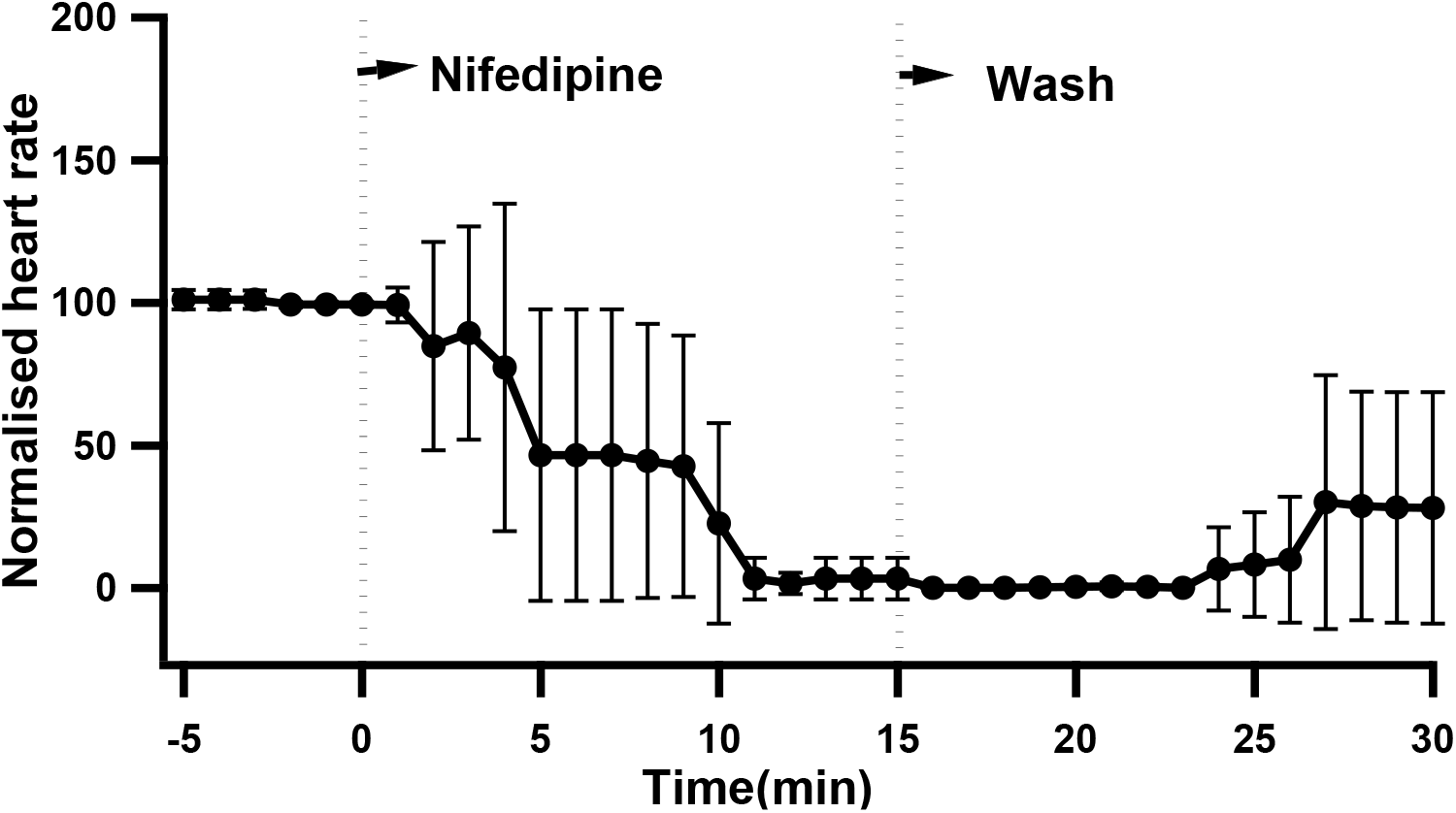
Nifedipine (10μM/L) solution stopped the isolated perfused rat heart completely after 10 minutes of perfusion.

**Figure 7:**
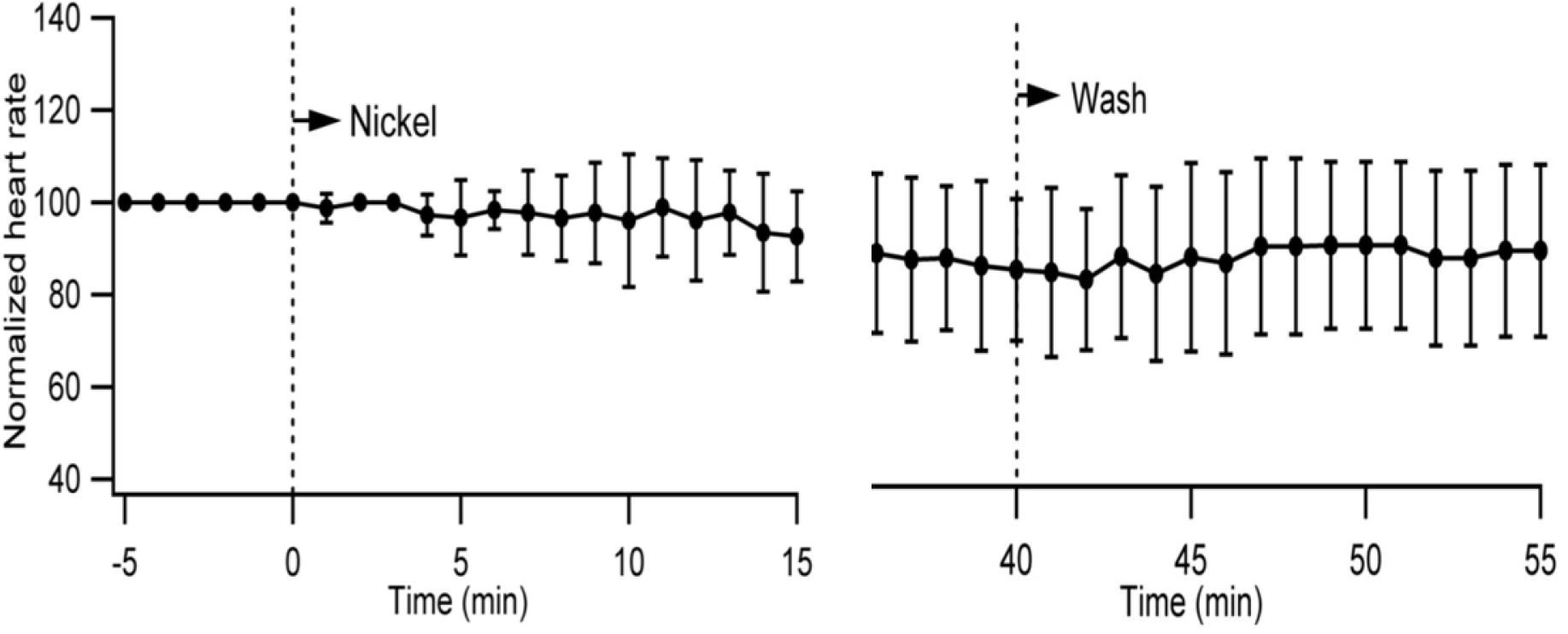
Nickel (40 μmol/L), did not have any effect on the rate of beating of the isolated rat heart even after 50 minutes of perfusion.

**Figure 8:**
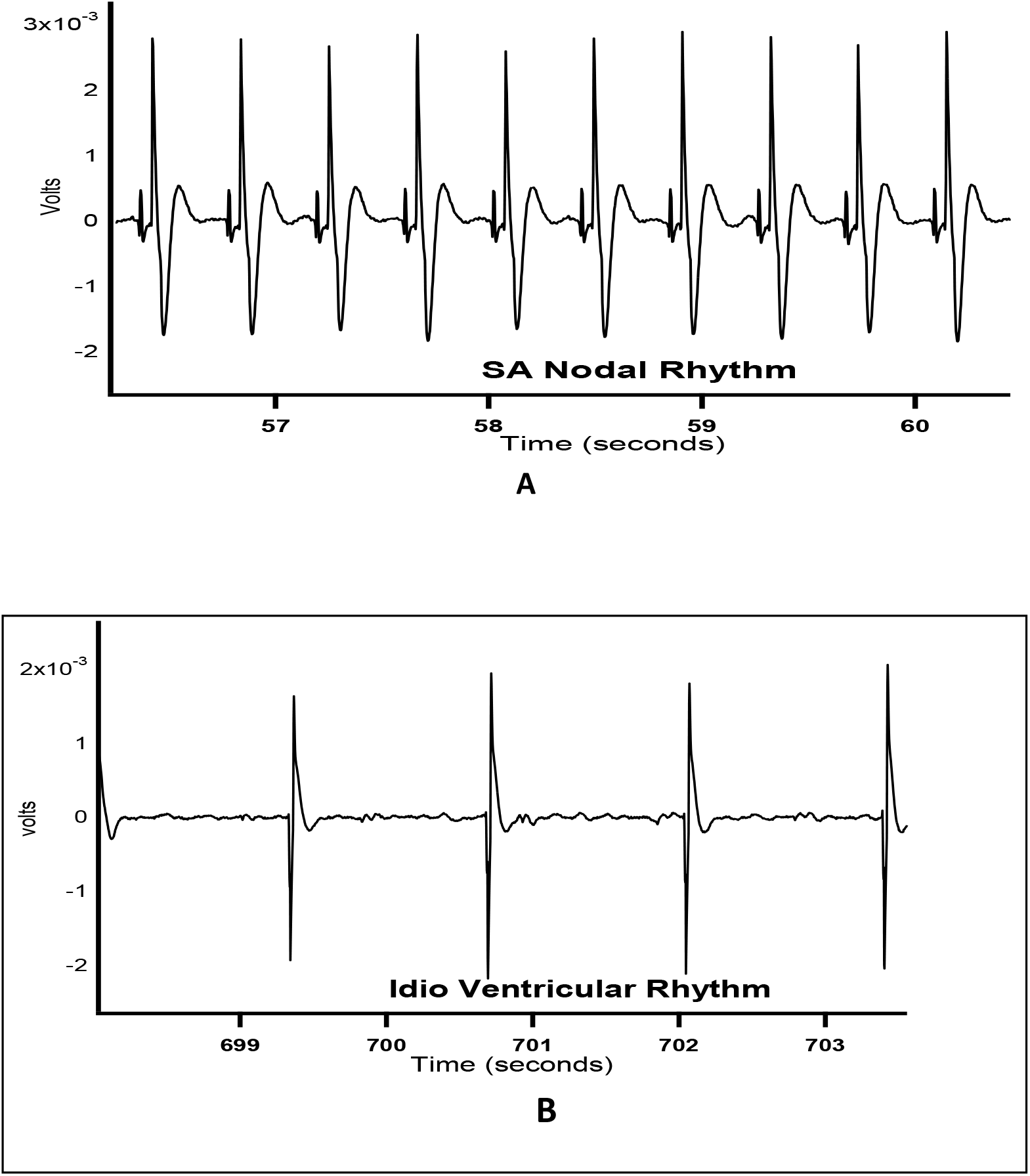
A: Surface electrogram from isolated perfused rat ventricle beating with SA nodal rhythm. P waves are seen. B: Surface electrogram from a rat heart beating at Idio-ventricular rhythm. P waves are absent.

### Effects of Nickel (40 μmol/L)

With Nickel (40 μmol/L), the heart rate changed from 132 ± 15 to 115 ± 36 bpm, showing only a 14.08 %reduction (p value = 0.206, n = 6). The reduction was not statistically significant. There was no change even after 50 minutes of perfusion with Nickel.

### Experiments on hearts with Idio-ventricular rhythm

On carefully cutting off the right atrium and severing the atrio-ventricular bundle by incising the inter-ventricular septum of the mounted and perfused heart, the heart stopped and then started to beat on its own (without any pacing). The new heart rate was low compared to the initial SA nodal rhythm and was considered as the idio-ventricular rhythm (Benforado, 1958). The basal heart rate in ventricular rhythm was 57 ± 16 (n= 21). The surface ECG had no P wave and the QRS complex was wide.

### Effects of Ivabradine on idio-ventricular rhythm

In 4 out of 6 experiments, ivabradine reduced the idio-ventricular rate. In 2 preparations there was no change. On averaging from all 6 experiments, the heart rate changed from 58 ±21 to 25 ±20 bpm (p = 0.023, n = 5), The percentage reduction in idio-ventricular rate was 58 ±31% (Figure 9).

**Figure 9:**
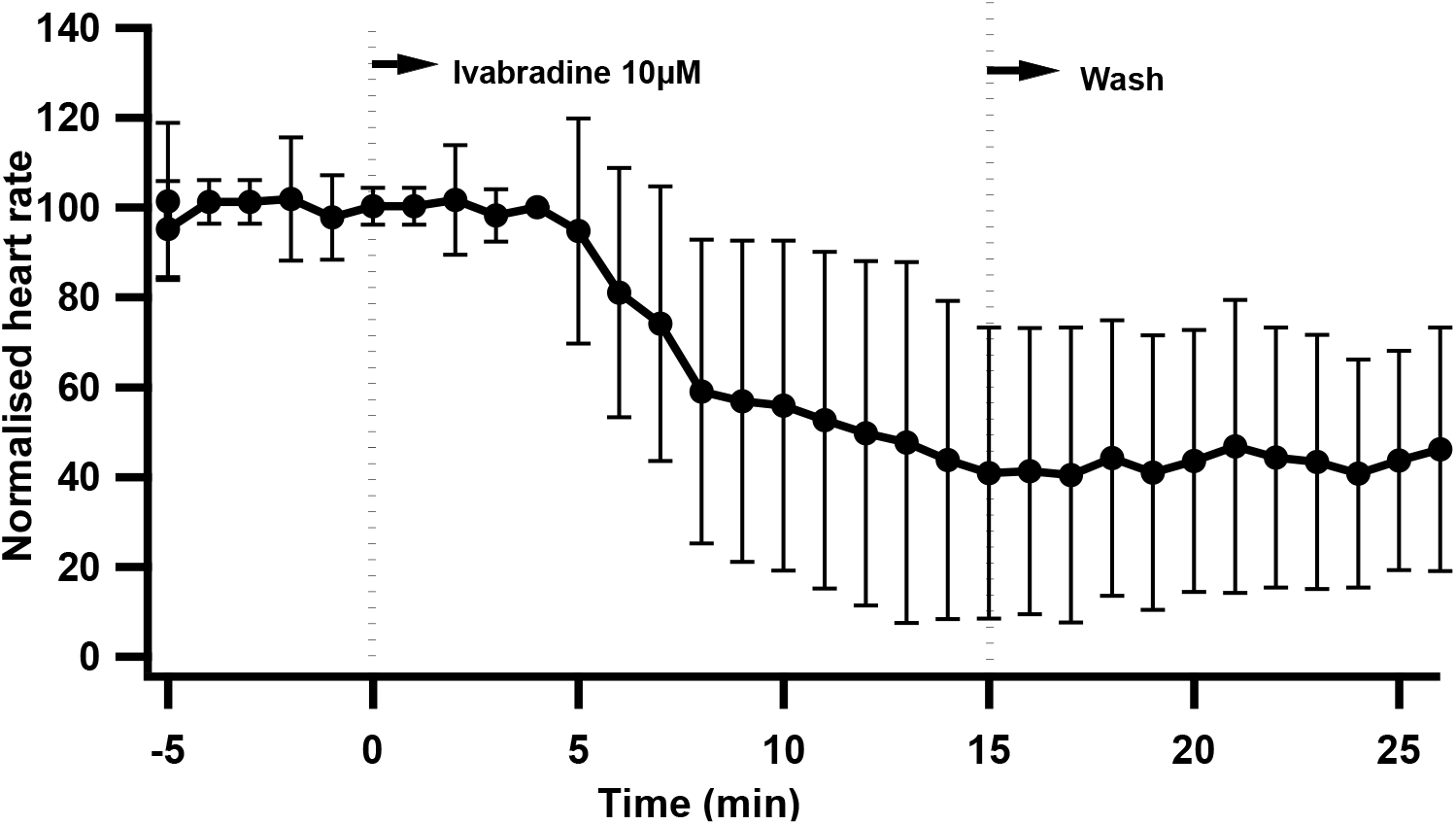
Ivabradine 10μM reduced the heart rate in isolated perfused rat heart in idio-ventricular rhythm but did not stop the heart.

### Effects of Low sodium on idio-ventricular rhythm

Isolated hearts in idio-ventricular rhythm with a basal heart rate of 58 ± 6 bpm in PSS, progressively slowed down and stopped beating within 5 minutes (p value = 0.043, n=5), upon switching to a low sodium solution. After sodium replacement by perfusing with PSS, heartbeat slowly recovered (26 ± 12 bpm) but did not reach pre-intervention values (Figure 10).

**Figure 10:**
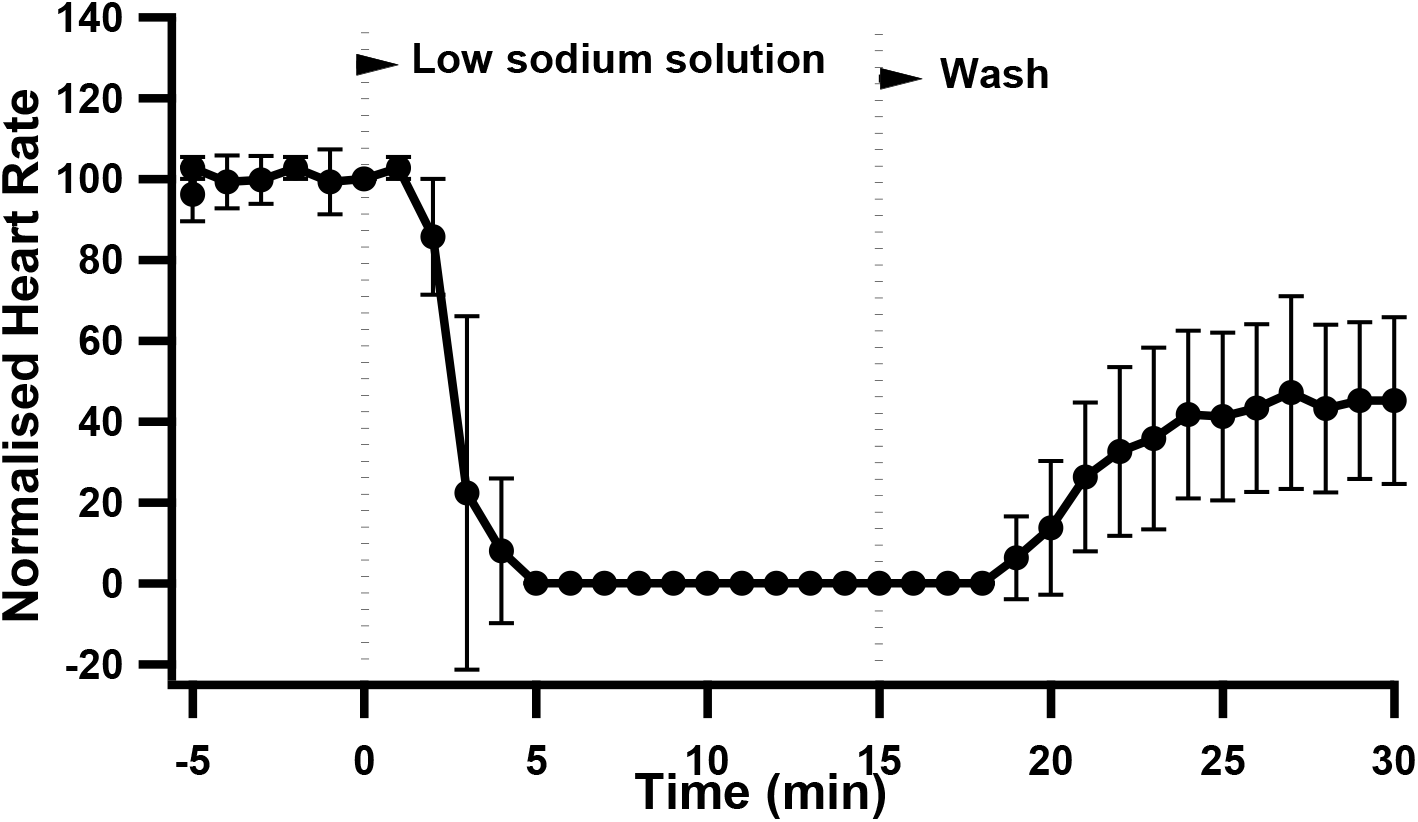
Low sodium solution (35 mmol/L sodium) stopped the isolated rat heart in idio-ventricular rhythm within 5 minutes of perfusion.

### Effects of Nickel on idio-ventricular rhythm

With 40μM Nickel which blocks the T-type calcium channels, there was no significant change in idio-ventricular heart rate. The heart rate before intervention was 53 ± 25 bpm which became 58 ±29 bpm after 15 minutes of perfusion with Nickel (Figure 11). The change was not statistically significant (p = 0.5, n= 5).

**Figure 11:**
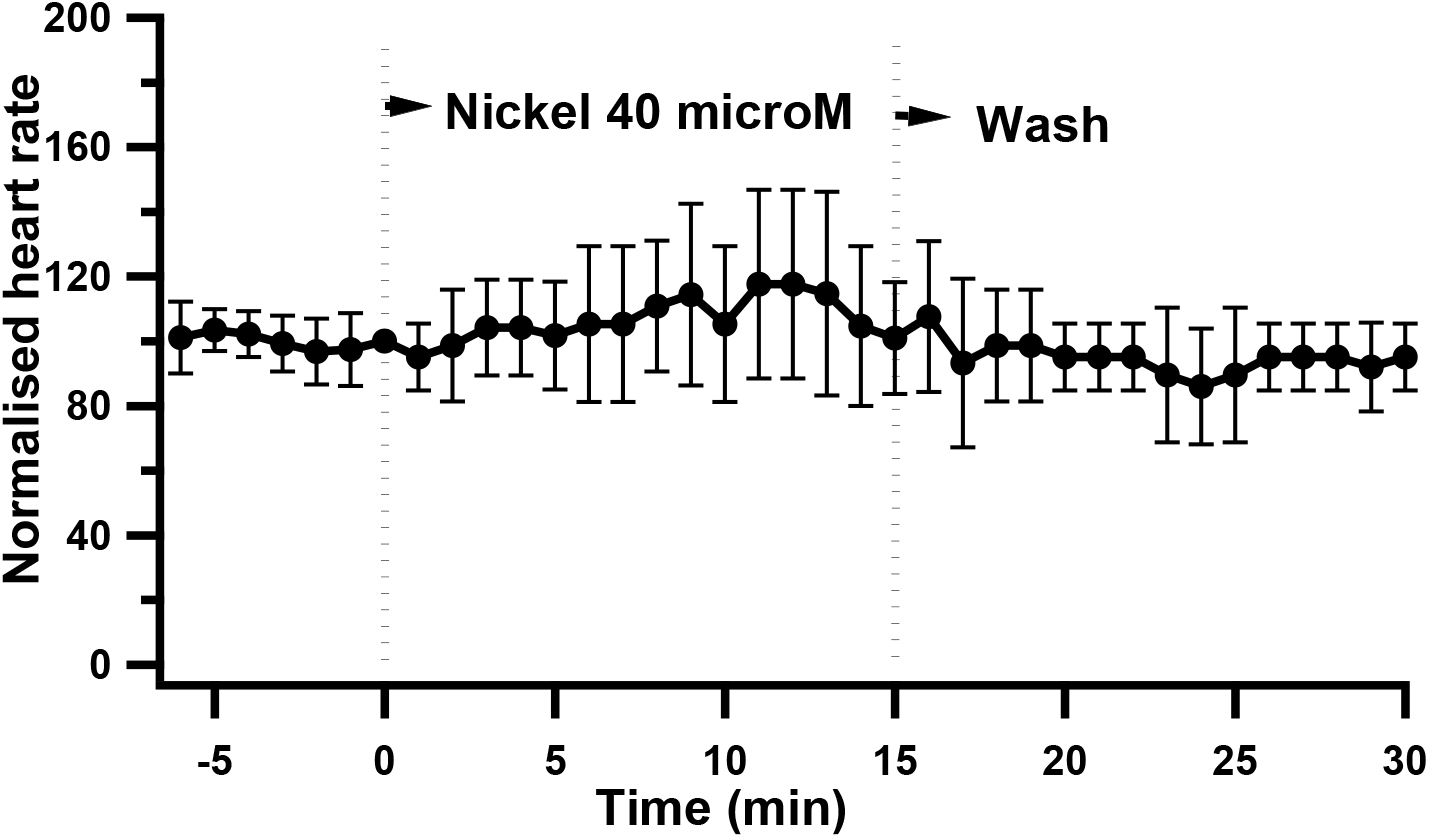
Nickel (40 μmol/L), did not have any effect on the heart rate of the isolated rat heart in idio-ventricular rhythm.

### Effect of Nifedipine on idio-ventricular rhythm

10 μmol/L Nifedipine, produced varied responses. It stopped the idio-ventricular rhythm only in 2 out of 5 preparations (Fig.12. A, B). Nifedipine increased the rate in 2 preparations (Fig.12.C, D). In one of these the rate continued to remain high even after wash (Fig.12. C), and in the other, the higher rate was maintained as long as Nifedipine was perfused. On washing Nifedipine with PSS, the heart stopped followed by recovery to pre-intervention levels (Fig.12. D). The heart rate did not change much with Nifedipine in one out of 5 preparations. (Fig.12.E).

**Figure 12:**
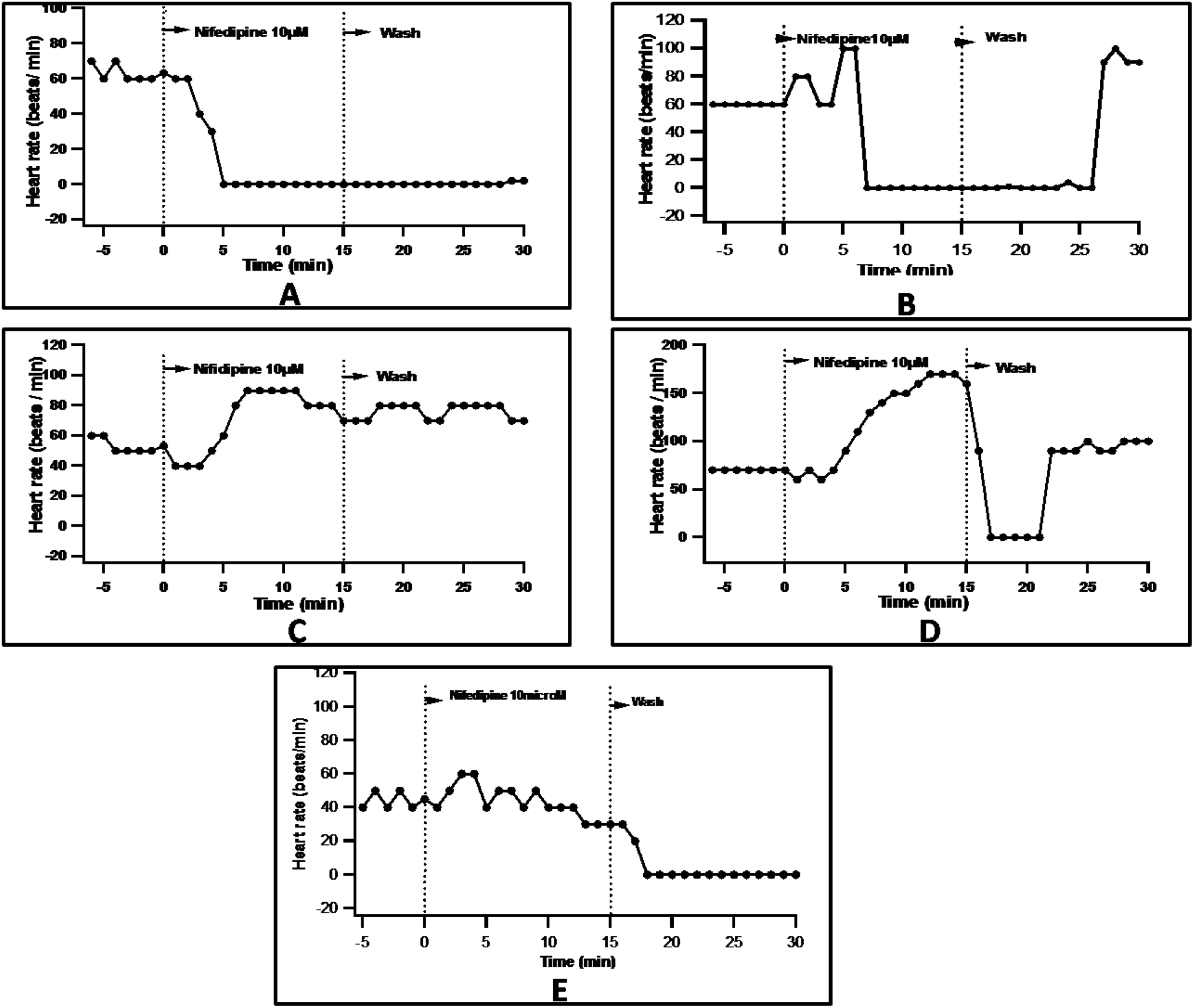
A-E: 10 μmol/L Nifedipine, produced varied responses in 5 different experiments.

## Discussion

In this study, Cesium, a known non-specific extracellular blocker of If was used at a dose of 5 mmol/L to block it (Denyer and Brown, 1990). It reduced the heart rate in the intact whole heart (beating at a rhythm set by the SA node) by 26% but did not stop the heart. The effect of cesium is not entirely due to blockade of If, because it also can block potassium channels, and hence favour depolarization and an increase in heart rate. What is observed probably is a net effect of blockade of If and inhibition of potassium channels, the two effects opposing each other.

Ivabradine blocks If completely at a dose of 3μmol/L (Ceconi et al., 2009). However, on the intact isolated heart, even at a dose of 10 μmol/L, it did not stop the heart completely but caused a mean reduction of heart rate by 46%. Similarly, Ivabradine produced 58 % percent reduction in ventricular rate, but did not stop the idio-ventricular rhythm. Since heart rate reduces by about half when If is inhibited, the inference is that it is an important rhythm-generating mechanism. However, it is not mandatory for SA node rhythm or idio-ventricular rhythm in the rat heart. The issue of whether If is the primary pacemaker current or not has been extensively debated (DiFrancesco & Noble 2012; Lakatta & Maltsev 2012).

Calcium-extrusive or forward mode I_NCX_, which would give rise to an inward current, was inhibited in the intact isolated heart by two different interventions namely: (1) low sodium solution (2) sodium-free solution where lithium was used as a substitute for sodium. Both these interventions abolished the electrical activity of the heart. Since what is essentially recorded on the electrogram is ventricular electrical activity, and since upstroke of the ventricular action potential is due to a sodium current, it had to be ensured that removal of sodium to block NCX does not impair the ability of the ventricle to develop action potentials. Replacement of up to 75% of sodium with an impermeant ion like choline does not reduce the peak of the action potential, although the rate of rise is less. If external sodium is replaced with lithium which can permeate through voltage-gated sodium channels, even the rate of rise of action potential is not affected (Niedergerke and Orkand, 1966). The complete arrest of SA nodal rhythm in the low sodium or sodium-free lithium solutions is therefore, not due to an inability of the ventricle to develop phase 0 of the ventricular action potential in response to an impulse, but due to an inability of the SA node to generate an impulse. In the hearts beating with an idio-ventricular rhythm too, NCX reversal with the low sodium solution stopped the heart. The inference is that INCX in forward mode is mandatory for rhythm generation in rat SA node and ventricle.

The L-type Ca^2+^ channel blocker Nifedipine (10 μmol/L), abolished the electrical activity of the SA node. It is understandable that I_CaL_ is mandatory for SA node rhythm generation because the upstroke of SA node action potential is due to I_CaL_. It is not possible to conclude from this experiment, whether I_CaL_ has a role to play in phase 4 diastolic depolarization in the SA node. On the other hand, Nifedipine did not abolish the idio-ventricular rhythm in some hearts and even increased the idio-ventricular rate in some. The lack of cardiac arrest with Nifedipine in some experiments allows us to infer safely that I_CaL_ is not mandatory for the idioventricular rhythm directly. However, since it is the source of calcium replenishment for the SR, block of I_CaL_ may result in depletion of SR in the longer run, and therefore abolition of LCR, which is the driver for forward mode I_NCX_. Such non-replenishment of SR calcium due to Nifedipine can account for the arrest of rhythm seen in some experiments with Nifedipine. The increase of heart rate with Nifedipine in some experiments is surprising. Nifedipine has also been shown to accelerate ventricular tachycardia in dogs (Brachmann *et al*., 1989). Since I_CaL_ occurs during the plateau phase, blocking it with Nifedipine will shorten the duration of the plateau. Shortening of action potential duration (APD) with nifedipine has been documented (Amlie *et al*., 1979, Go *et al*., 2005). While the dependence of APD on heart rate is well-documented (Li *et al*., 1999, Arnold *et al* 1982), it remains to be seen if the converse is true. Is heart-rate dependent on APD? It seems only logical that a reduction in APD and therefore wavelength should increase frequency. If reduction in APD can increase ventricular rate (as in the case with Nifedipine), and if I_NCX_ mediated calcium efflux is a mandatory rhythm-generating mechanism as is suggested by our results, then it follows that the probability of LCR responsible for driving I_NCX_ in forward mode is increased by repolarization. Some studies have ruled out the dependence of LCR on cyclical changes on membrane depolarization (Maltsev and Lakatta, 2007) or on calcium influx (Vinogradova et al., 2002), However, Huser *et al* (2000) have shown that LCR occurs at negative potentials of about – 60 mV. Tsutsui *et al* (2018) also report that while LCR decreases as the membrane potential reaches the maximum diastolic potential (MDP), it begins to increase soon after MDP, and an action potential ensued. Therefore, it is logical to conclude that the increase in heart rate seen with nifedipine in the ventricular preparation is due to the fact that nifedipine-induced shortening of APD caused LCR to happen quicker than normal, and therefore initiated I_NCX_, earlier than normal.

Increasing external calcium can have two opposite effects on membrane potential. It can either reverse NCX and therefore inhibit depolarization or enhance I_CaL_ and therefore enhance depolarization. The effect on heart rate would depend on which of the above two effects is dominant. The results obtained with high calcium solution on the SA node corroborate such a view. There was an increase in heart rate in half the number of experiments and a decrease in the other half.

Studies on low-voltage-gated T type Ca^2+^ channels have shown that they can be selectively blocked by 40 μmol/L Nickel (Hagiwara et al., 1988; Huser et al., 2000). Though in cat atrial pacemaker cells Nickel reduced the rate of spontaneously beating cells (Huser et al., 2000), in rabbit SA node cells, diastolic calcium release is not affected by block of I_CaT_ (Vinogradova et al., 2002). In the present study too, block of I_CaT_ with nickel did not change either the SA node rate or the idio-ventricular rate. Hence it is inferred that I_CaT_ does not play a significant role in rhythm generation in the SA node or ventricle.

To summarize, I_NCX_ and I_CaL_ are obligatory for rhythm generation in the rat SA node while I_NCX_ is the only obligatory current for idio-ventricular rhythm. If contributes significantly to rhythm generation in both SA node and ventricle but is not mandatory. Heart rate is reduced by about half (SA node rhythm) or more than half (idioventricular rhythm) with If blockade. I_CaT_ does not play a role in either rhythm. Since the idio-ventricular rate is abolished only on reversal of I_NCX_ and not by blockade of other currents, it may be argued that the diastolic calcium release (LCR) responsible for driving I_NCX_ occurs independent of preceding membrane events. However, frequency of LCR is doubled or more than doubled by If. Our conclusion therefore is that If and I_NCX_ contribute equally to pace-making in both sinoatrial node and the ventricle in the rat heart. This is in corroboration of the coupled-clock pace-maker theory proposed by Maltsev and Lakatta (2012).

## Notes

### Competing Interest Statement

The authors have declared no competing interest.

